# An organic extract from ascidian *Ciona robusta* induces cytotoxic autophagy in human malignant cell lines

**DOI:** 10.1101/2023.11.18.567676

**Authors:** Alessandra Gallo, Ylenia Maria Penna, Maria Russo, Marco Rosapane, Elisabetta Tosti, Gian Luigi Russo

**Affiliations:** Department of Biology and Evolution of Marine Organisms, Stazione Zoologica Anton Dohrn, Villa Comunale, 80121, Naples, Italy; National Research Council, Institute of Food Sciences, 83100, Avellino, Italy

**Keywords:** Marine organisms, ascidians, Ciona *robusta*, cytotoxicity, autophagy, cell viability, anticancer agents

## Abstract

The last decades have seen an increase in the isolation and characterization of anticancer compounds derived from marine organisms, especially invertebrates, and their use in clinical trials. In this regard, ascidians, which are included in the subphylum Tunicata, represent successful examples with two drugs, Aplidine© and Yondelis©, that reached the market as orphan drugs against several malignancies. Here, we report that an organic extract prepared from homogenized tissues of the Mediterranean ascidian *Ciona robusta* inhibited cell proliferation in HT-29, Hep G2, and U2 OS human cells with the former resulting as the most sensitive to the extract (EC_50_ = 250 µg/ml). We demonstrated that the ascidian organic extract was not cytotoxic on HT-29 cells induced to differentiate with sodium butyrate, suggesting a preference for the mixture for the malignant phenotype. Finally, we reported that the cell death induced by the organic extract was mediated by the activation of a process of cytotoxic autophagy as a result of the increased expression of the LC3-II marker and the number of autophagic vacuoles, which almost doubled in treated HT-29 cells. In summary, although the detailed chemical composition of the *Ciona robusta* extract is still undetermined, our data suggest the presence in it of bioactive compounds possessing anticancer activity.

## 1 Introduction

The last decades have seen increased isolation and characterization of anticancer compounds derived from marine organisms and their use in clinical trials. The initial studies in this field, dating back to the seventies of the last century, originated from the interest of naturalists and marine biologists, who identified various toxins present only in the organisms that inhabit marine ecosystems. Examples are Conidia, a family of marine gastropod mollusks that inject a potent peptide toxin (conotoxin) to immobilize prey (McManus et al., 1981); the cnidarians Zoanthids, commonly found in coral reefs, possessing a toxic polyketide, “palytoxin”, which makes them unpleasant to predators (Moore and Scheuer, 1971); some microalgae producing a highly cytotoxic alkaloid neurotoxin, saxitoxin, and a polyketide neurotoxin, brevetoxin (Simmons et al., 2005). In recent years, the studies to isolate new substances with anticancer activity have intensified thanks to a series of research programs supported by various institutions, including the National Cooperative Drug Discovery Program of the National Cancer Institute (NCI). Subsequently, the development of collaborations between groups of academic researchers and major pharmaceutical companies, including NCI, led to the development of several agents that entered clinical trials, excellently recently reviewed in recent works (Sorokina and Steinbeck, 2020; Cooreman et al., 2023). In these studies, interesting links can be found to updated databases on natural products including those of marine origin.

Although the main marine sources of bioactive compounds are represented by invertebrates, largely sponges, tunicates and soft corals, the observation that microorganisms such as marine microalgae, cyanobacteria and heterotrophic bacteria live in symbiosis with them led in the late nineties to formulate the “symbiotic origin” hypothesis for some of the most bioactive and druggable natural products (Piel, 2004; 2009) (Haygood et al., 1999; Piel, 2004; 2009). Metagenomics analyses have validated this concept, providing insights on the possibility of culturing marine microorganisms as resources for new classes of therapeutic drugs (Williams, 2009; Cooreman et al., 2023).

Successful examples come from marine species of the class Ascidiacea included in the subphylum Tunicata, phylum Chordata. Ascidiacea, commonly known as ascidians or sea squirts, are grouped into three orders: Aplousobranchia, Phlebobranchia and Stolidobranchia. The genomes of twenty ascidian species are deposited in the NCBI databanks^1^. Plitidepsin (dehydrodidemnin B; Aplidine©) is a cyclic depsipeptide with anticancer properties originally isolated from Caribbean tunicate *Trididemnum sp*. (Rinehart et al., 1981). Subsequently, the same compound was isolated from the marine α-proteobacterium *Tistrella mobilis*, hosted by the same tunicate (Tsukimoto et al., 2011). Plitidepsin, under the commercial name of Aplidine©, has been approved as an orphan drug for several hematological malignancies, such as acute lymphoblastic leukemia, multiple myeloma, primary myelofibrosis, post essential thrombocythaemia myelofibrosis (Alonso-Álvarez et al., 2017; Cooreman et al., 2023). More recently, promising preclinical effects have been demonstrated for Aplidine© in treating SARS-CoV-2 infection in two different mouse models suggesting potential therapeutic applications against COVID-19 (White et al., 2021).

More than 35 years have passed since the discovery of another bioactive molecule, which was originally listed among the most promising antibiotic anticancer drugs derived from marine organisms, Ecteinascidin-743 (also named ET-743, trabectedin, Yondelis©) (Rinehart et al., 1990). ET-743 was isolated from the tunicate *Ecteinascidia turbinate* and later identified as a product of the bacterial symbiont *Candidatus Endoecteinascidia frumentensis* (Rath et al., 2011). The mechanism of ET-743 is unique in that it binds to the DNA minor groove and alkylates guanine in the N2 position preferentially after a GG or GC sequence. These ET-743-DNA adducts are recognized by the NER (DNA - Nucleotide Excision Repair) repair system but unlike other adducts caused by alkylating agents such as cisplatin, the NER system causes not DNA repair but cell death even at very low concentrations (10-100 ng/ml) and preferentially via the mitochondrial pathway, caspase-3 and JNK kinase activation (Takebayashi et al., 2001; D’Incalci et al., 2002; D’Incalci and Galmarini, 2010). The clinical story of ET-743 is partially controversial. Between 2001 and 2003, under the name of Yondelis©, the drug received the status of orphan designation by EMA in the European Union for the treatment of soft tissue sarcoma and ovarian cancer. In 2015, the FDA approved it for treating soft tissue sarcoma, including unresectable liposarcoma and leiomyosarcoma in metastatic patients.

However, the conclusions reached by several trials on the toxicity and efficacy/safety balance of Yondelis© led the EMA and FDA agencies to re-evaluate the marketing authorization of Yondelis© (reviewed in (Cooreman et al., 2023)). Despite the examples of the symbiotic origin of plitidepsin and trabectedin, the synthetic source of the large part of active metabolites isolated from ascidians remains unknown (Tianero et al., 2015) and cannot be excluded that some of them may derive from unidentified dietary sources.

Considering the success of ascidian-derived agents in clinical trials, several years ago, searching for alternative sources of anticancer agents, we demonstrated that a methanolic extract prepared from the tissues of the ascidian *Ciona intestinalis* (class Ascidiacea order Phlebobranchia; family Cionidae) possessed antiproliferative and pro-apoptotic effect on cancer cell lines of different origin (Russo et al., 2008). *C. intestinalis* represented an ideal candidate since its abundant presence in the Mediterranean Sea, its genome was deposited (De Luca Di Roseto et al., 2002; Satoh et al., 2003; Shi et al., 2005), and the anatomy, physiology and development were well known (Passamaneck and Di Gregorio, 2005; Tosti et al., 2011). However, it is important to clarify that the mentioned work on the methanolic extract from *C. intestinalis* (Russo et al., 2008) has been *bona fide* obtained from the specie *Ciona robusta*, not *C. intestinalis*. In fact, until 2015, *C. robusta* and *C. intestinalis* were formerly considered the same organism, being almost indistinguishable. Recent morphological, ecological, and genomic data reached the conclusion that the populations of *C. intestinalis* from the Mediterranean Sea, the Pacific Ocean (Australia, Japan, New Zealand, South Korea, and West coast of North America), and the Atlantic coasts of South Africa (formerly known as *C. intestinalis* Type A specimen) were similar to each other, but differed from the type B specimen of *C. intestinalis* present on the East Coast of North America, the coast of Northern Europe and the Bohai and Yellow Seas. Based on morphological and genetic data, it has been recently established that *C. intestinalis* Type A and *C. robusta* correspond (Caputi et al., 2007; Brunetti et al., 2015).

In the present work, we established a new protocol to isolate an organic fraction from homogenized tissues of *C. robusta* possessing antiproliferative activity on malignant cell lines. The bioactive extract was able to activate the process of cytotoxic autophagy.

## 2 Materials and methods

### 2.1 Chemicals

Crystal violet, formalin, chloroquine, sodium butyrate (NaBt), dimethyl-sulfoxide (DMSO), pNPP (p-nitrophenyl phosphate), Trypan blue were purchased from Merck/Sigma (Milan, Italy); gentamicin and kanamycin were purchased from Panreac Applichem (Darmstadt, Germany); PBS (phosphate-buffered saline) tablets were purchased from Euroclone (Milan, Italy). All other chemicals used were of research highest purity grade.

### 2.2 Preparation of Bioactive Extracts from ascidian *Ciona robusta*

This study did not include endangered or protected ascidian species and was conducted according to the guidelines of the Declaration of Helsinki amended by the European Directive 2010/63 on the protection of animals used for scientific purposes, transposed into the Italian law by Legislative Decree (2014)/26.

Bioactive extracts from *C. robusta* were prepared by combining different protocols previously published (Sakai et al., 1992; Russo et al., 2008) and summarized in the scheme reported in Figure 2. Briefly, adults of *C. robusta* were collected from different marine sites in the Mediterranean Sea (gulfs of Naples and Taranto, Italy), which are not privately owned or protected in any way, according to Italian legislation (DPR 1639/68, September 19, 1980, confirmed on January 10, 2000). After collection, ascidians were transported to the Marine Biological Resources Service of Stazione Zoologica Anton Dohrn and acclimated for at least 7 days before use in tanks (1 animal/L) with running natural seawater (temperature of 18 ± 2 °C, pH 8.1 ± 0.1, salinity 39 ± 0.5 psu) equipped with oxygen pumps and fed daily with the shellfish diet 1800.

Small groups of samples (3-4 individuals depending on their length ranging between 2-5 cm) were collected, their external tunic removed and the seawater in excess eliminated by gently squeezing. The remaining tissues were transferred in a 50 ml plastic tube, chopped in small pieces with scissors to facilitate the homogenization and added with about three volumes (w/v) of *i*-PrOH. The crude homogenate obtained after homogenization using an Ultra-Turrax T25 blender (20,000 rpm; 5 min) was centrifuged at 3000xg for 10 min. The *i*-PrOH extract was concentrated by a rotovapor to obtain an aqueous emulsion, which was extracted three times in an equal volume of ethyl acetate using a pear-shaped separatory funnel. The three organic, upper phases (OrPh) were combined, dried by a rotovapor, and then redissolved in MeOH. The material soluble in MeOH was quantitate by weight and a stock solution of 25 mg/ml was prepared for the subsequents cytotoxicity assays. An aliquot of the lower aqueous phase (AqPh), which contained about 90% in weight of the total material present in the aqueous emulsion after the *i*-PrOH extraction was brought to a stock solution of 200 mg/ml to be applied to cell cultures. Both the organic and aqueous phases were stored at −20°C.

For the experiments in the presence of antibiotics, fifteen individuals of *C. robusta* were reared for three weeks in a closed system tank containing natural seawater and a mixture of gentamicin and kanamycin (100 mg/L) at the same condition of acclimatization. The seawater was changed and antibiotic solutions were prepared daily. Control individuals were maintained in identical experimental conditions avoiding the addition of antibiotics.

### 2.3 Cell Culture and Reagents

The HT29 cell line, derived from a forty-four-year-old Caucasian patient suffering from colorectal adenocarcinoma (Von Kleist et al., 1975) was purchased from the American Tissue Culture Collection (ATCC), LGC Standards (Sesto San Giovanni, Milan, Italy). The Hep G2 cell line, isolated from a hepatocellular carcinoma of a 15-year-old male youth with liver cancer (Aden et al., 1979) and the U2 OS cell line, derived from a moderately differentiated sarcoma of the tibia of a 15-year-old female osteosarcoma patient (Ponten and Saksela, 1967) were kindly donated by prof. M.A. Belisario from University of Salerno (Salerno, Italy) and prof. A. Oliva from L. Vanvitelli University (Naples, Italy), respectively.

Cell lines were cultured at 37°C in a humidified atmosphere with 5% CO2 using Dulbecco’s Modified Eagle’s Medium (DMEM; EuroClone, Pero, Milan, Italy) supplemented with 10% foetal bovine serum (FBS; EuroClone) with 100 µg/ml penicillin/streptomycin, 2 mM L-glutamine, and 100 µM non-essential amino acids (EuroClone). To avoid any variations caused by the long-term culture of the cells, we used early passages (<10) to ensure reproducibility. Cell propagation was carried out by treating them with a trypsin/EDTA solution (EuroClone) when confluence was reached and counting/replating using Trypan blue exclusion dye in an automatic cell counter (EveTM, NanoEnTek distributed by VWR, Milan, Italy) in accordance with the manufacturer’s instructions.

Treatments with the *C. robusta* extracts was performed at the times and concentrations indicated in the Results and Discussion section and in the figure legends. At the end of the experiments, cells were analyzed to measure changes in cell viability, autophagy and expression of autophagy markers as reported below. Cells were plated at an appropriate density in quadruplicate in a 96-well plate and allowed to adhere for 24 h before performing the functional assays.

Differentiation of HT-29 cells was obtained by treatment for 0-120 h with 4 mM NaBt as described (Russo et al., 1997; Russo et al., 1999) before the addition of the *C. robusta* extracts.

We measured the formation of H_2_O_2_ by FOX (ferrous oxidation-xylenol orange) assay as reported (DeLong et al., 2002) after incubation of *C. robusta* extracts at the concentrations indicated in the Results and Discussion with cell culture medium in the absence of cells. The FOX assay is based on the oxidation of the ferrous (II) ion to the ferric (III) ion in an acid environment. The molecular complex formed by the ferrous ion (II) and the relative is determined calorimetrically. A specific application of the fox method is to identify oxidant species produced in the cell culture medium by the pro-oxidant activity of antioxidant molecules or extracts in the presence of high concentrations of metal ions (Fenton reaction).

### 2.4 Cell Viability Assays

Cell viability assay was performed primarily using the Crystal Violet technique, which uses a protein-binding dye (Tedesco et al., 2013; Feoktistova et al., 2016). Briefly, to eliminate any residue of the culture medium, before staining a washing was carried out for 15 min with phosphate buffer saline solution (PBS) followed by the addition of 10% formalin in PBS to allow fixation. Finally, Crystal Violet 0.02% (w/v) in an aqueous solution was incubated for 30 min before dye removal and observation of the stained cells in a bright field invertoscope (Axiovert 200, Zeiss) with 200-400X magnification to allow high-resolution photographs to be taken. Quantification was carried out by adding a 10% (v/v) acetic acid solution to solubilize the dye incorporated within the cells followed by a spectrophotometric reading at a wavelength of 595 nm using a microplate reader (Synergy HT BioTek, Milan, Italy).

CyQuant (Thermo-Fisher Scientific/Life Technologies, Milan, Italy) method was performed essentially as previously described (Russo et al., 2017b). In the presence of the rea-gent called “background suppressor” which penetrates through the pores that form on the membranes of damaged cells, the CyQuant dye can no longer intercalate into the nuclear DNA, and the dead cells do not fluoresce. Briefly, CyQuant mixture was added to the cells following incubation for 1 h at 37°C following the manufacturer’s instructions. The CyQuant kit contains the nuclear dye (CyQuant nuclear stain; 1:250 dilution) and the suppressor of basal fluorescence (background suppressor; 1:50 dilution), which discriminates between living and dead cells, avoiding the staining of the latter. Fluorescence was measured at the excitation wavelength of 485 nm and 530 nm emission. The results were expressed as a percentage of fluorescence of the untreated control using a microplate reader (Synergy HT BioTek).

### 2.5 Measurement of Autophagy

Autophagy was monitored in live cells using the Cyto-ID Autophagy Detection kit (Enzo Life Sciences, Milan, Italy) (Russo et al., 2023). Autophagy activation was detected using this specific kit based on a fluorescent amphipathic cationic tracer able to specifically detect the number of intracellular autophagosomes and not in the lysosomes. Cells (approx. 1×10^4^ cells per well) were plated in a 96-wells plate (Corning-Costar, EuroClone) and treated with the *C. robusta* extracts as indicated in the Results and Discussion section. After treatment, the medium was removed and the cells washed with Assay Buffer containing 5% FBS. The mixture (100 µL) with the autophagy detection marker (Cyto-ID, diluted 1:500) and nuclear dye (Hoechst 33342, diluted 1:1000) was added in phenol red-free DMEM at 5% FBS and allowed to react for 30 min in the incubator at 37°C in 5% CO_2_ atmosphere. The cells were washed with Assay Buffer before being photographed and determining a double fluorometric measurement using a microplate fluorescence reader (Synergy HT BioTek). The autophagosomes were quantified by normalizing green fluorescence (Cyto-ID) at an excitation wavelength of 495 nm and an emission wavelength of 519 nm, as for FITC (fluorescein isothiocyanate). The second measurement concerning the blue fluorescence (Hoechst dye 33342) was performed at the wave-lengths of 358/461 nm as for DAPI (4’,6-diamidin-2-phenylindole). The FITC/DAPI ratio obtained in fluorescence units allowed the quantification of cells with autophagosomes compared to the total number of cells. A fluorescence microscope invertoscope (Zeiss Axiovert 200; 400x magnification) was used to observe and take photos of autophagic vesicles and nuclei through the application of FITC and DAPI filters.

### 2.6 Polyacrylamide Gel Electrophoresis and Immunoblotting

HT-29 cells (generally 1.5×10^6^) treated with *C. robusta* extracts were lysed using a lysis buffer containing protease and phosphatase inhibitors (50 mM Tris/HCl, pH 7.4; 150 mM NaCl; 5 mM ethylenediaminetetraacetic acid; 1% Nonidet P-40; 0.5 mM dithiotreitol; 1 mM Na_3_VO_4_; 40 mM NaF; 1 mM Na_4_P_2_O_7_; 7.4 mg/ml 4-p-nitrophenyl phosphate; 10% glycerol; 100 μg/ml phenylmethylsulfonyl fluoride and the cocktail of protease and phosphatase inhibitors (Merk/Sigma) (Russo et al., 2017a). After protein concentration determination using the Bradford method (Bradford, 1976), the total lysates (20-30 µg/lane) were added with a solution to reduce the disulfide bridges (Reducing Agent 20X; Bio-Rad, Milan, Italy) and Laemmli Sample Buffer 4X (Bio-Rad) (100 mM Tris/HCl pH 6.8; 4% SDS; 200 mM dithiotreitol; 20% glycerol; 0.2% bromophenol blue dye). The samples were first heated for 5 min at 95°C and then centrifuged at 11,000xg in microfuge before loading them on 4-12% pre-cast mini-or midi-gels (Thermo-Fisher Scientific/Life Technologies). The running buffer contained MOPS [3-(N-morpholino) propanesulfonic acid], (50 mM MOPS, 50 mM Tris, 1% SDS, 1 mM EDTA, pH 7) for the separation of high molecular weight proteins, while MES buffer [2-(N-morpholino) ethanesulfonic acid] (50 mM MES, 50 mM Tris, 1% SDS, 1 mM EDTA, pH 7) was used for low molecular weight proteins. A constant voltage (200 V) was set for the electrophoretic chamber (Thermo-Fisher Scientific/Life Technologies). At the end of electrophoresis, proteins were transferred from the gel to a 0.2 µm polyvinyldenfluoride (PVDF) membrane (Transfer pack; Bio-Rad) using the TransBlot Turbo system (Bio-Rad) applying a constant amperage (2.5 mA) for 7 min at room temperature. After transfer, gels were stained using Coomassie blue R-250 (Merck/Sigma) to detect the untransferred proteins, and the membranes washed with 1X T-TBS (Tween-20-Tris Buffered Saline) (0.1% Tween-20; 25 mM Tris, pH 8; 137 mM NaCl; 2.69 mM KCl) for 5 min with shaking and incubated for 1 h with a blocking solution (3% bovine serum albumin, or not-fat dry milk 5% in T-TBS containing 0.02% NaN3). The following primary antibodies were diluted 1:1000 in 3% BSA/T-TBS and incubated for about 16 h: anti LC3-I/II (cat. # 12741S) from Cell Signaling Technologies (distributed by Euro-Clone); anti-α-Tubulin (cat. # T9026; Merck/Sigma). Following washing with T-TBS, the membranes were incubated for 2 h with a secondary antibody linked to horseradish peroxidase (diluted 1:20,000 in T-TBS). The immunoblots were developed using the ECL PrimeWestern blotting detection system kit (GE Healthcare, Milan, Italy). Band intensities were quantified and expressed as optical density on a Gel Doc 2000 Apparatus (Bio-Rad) and Multianalyst software (Bio-Rad).

### 2.7 Statistical Analysis

The results have been expressed as mean ± standard deviation (SD) based on the values obtained from independent experiments carried out in duplicate, triplicate or quadruplicate. Differences between the two groups were analyzed using the Student’s t-test (Excel MS software 2016), and significance was established as indicated in the legends of the figures.

## 3 Results and Discussion

To verify the capacity of the *C. robusta* organic extract to affect cell viability in cancer cells, we selected three cell lines, namely HT-29, Hep G2, and U2 OS derived from a human colorectal adenocarcinoma cell line, a hepatocellular carcinoma, and a moderately differentiated osteosarcoma, respectively. These cell lines were selected based on the following criteria: i) they grow adherent and have in common an epithelial-like morphology even deriving from different cancer types; ii) they are generally resistant to cell death induced by chemotherapeutic and/or pro-apoptotic drugs; iii) they represent examples of tumors responding to the anticancer activity of other ascidian-derived agents (Cooreman et al., 2023).

As reported in Figure 1, the original isopropyl alcohol (*i*-PrOH) extract (Figure 2) showed a different grade of cytotoxicity on the selected cell lines, with HT-29 being the most sensitive with an EC_50_ of about 250 µg/ml compared to Hep G2 and U2 OS (EC50 > 500 µg/ml).

**Figure 1.**
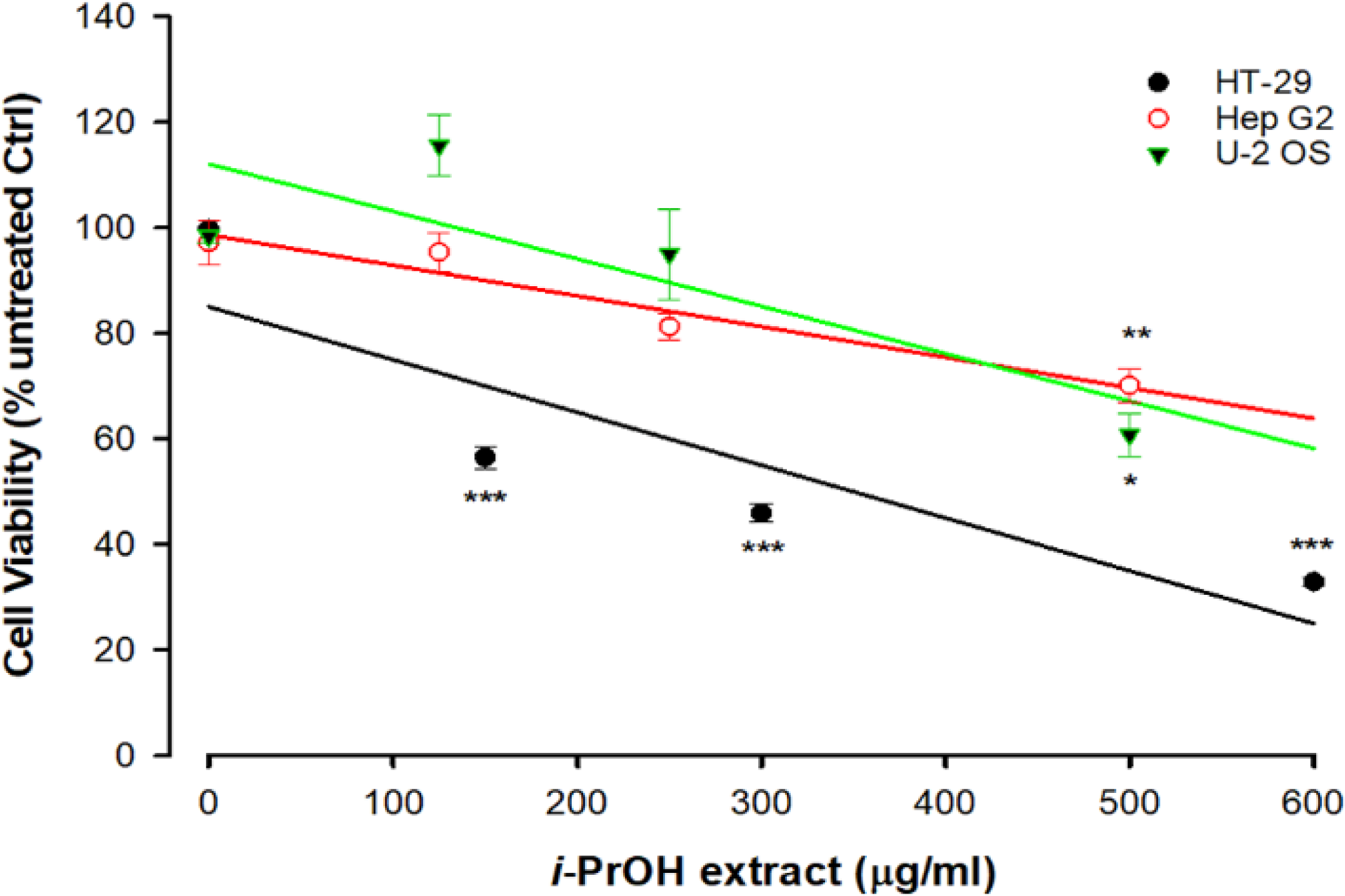
Cytotoxicity of *Ciona robusta* isopropyl extract on different cell lines. Cells (1×10^4^/well) were incubated with increasing concentrations of *i*-PrOH extract for 48 h. Cell viability was measured using the Crystal Violet assay, as reported in the Materials and Methods section. In the control (Ctrl) experiments (no *i*-PrOH extract), cells were treated with vehicle (methyl alcohol 2% v/v, final concentration). Data are presented as means of three independent experiments ± SD. Symbols indicate significance: ***P<0.0005; **P<0.005; *p<0.05 *vs* untreated Ctrl (Student’s t-test).

**Figure 2.**
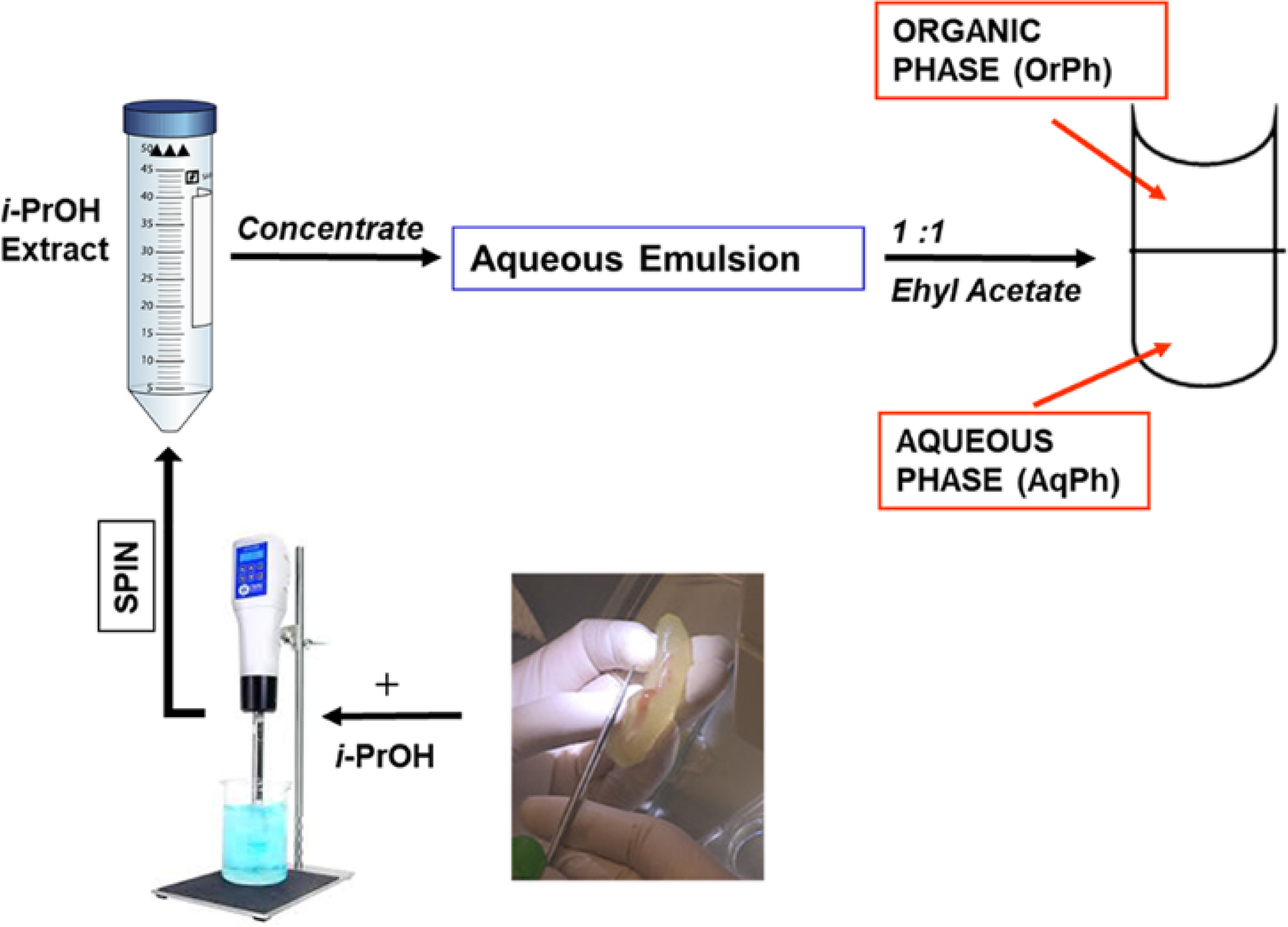
Preparation of the *Ciona robusta* extracts. The general scheme summarizes the different steps leading to the *C. robusta* extracts tested on the other cells lines reported in Figure 2. For details, see the description in the Materials and Methods section. *i*-PrOH, isopropyl alcohol; OrPh, organic phase; AqPh, aqueous phase.

Based on these results, HT-29 cells were selected as a leading cellular model for the next experiments. It is worthwhile to note that the screening of cell lines presented in Figure 1 extends the one previously reported (Russo et al., 2008) and indicates a selective cytotoxicity of the ascidian extract on specific cancer types with the human colorectal adenocarcinoma cell lines being among the most sensitive. In fact, HT-29 here and Caco-2 in our previous work (Russo et al., 2008), showed the lowest EC50.

The *i*-PrOH extract was further purified as indicated in Figure 2 and described in Materials and Methods to obtain two new phases, the aqueous (AqPh) and organic (OrPh) phases whose effects on cell viability were measured on the HT-29 cell line. Figure 3 shows that the AqPh was not associated with any cytotoxic activity. On the opposite, the OrPh was enriched in the agent(s) responsible for reducing cell viability resulting time-and dose-dependent. The calculated EC_50_ for the OrPh at 48 h resulted in about 90 µg/ml, three times lower than the value reported in Figure 1 for HT-29 cells and referring to the crude *i*-PrOH extract.

**Figure 3.**
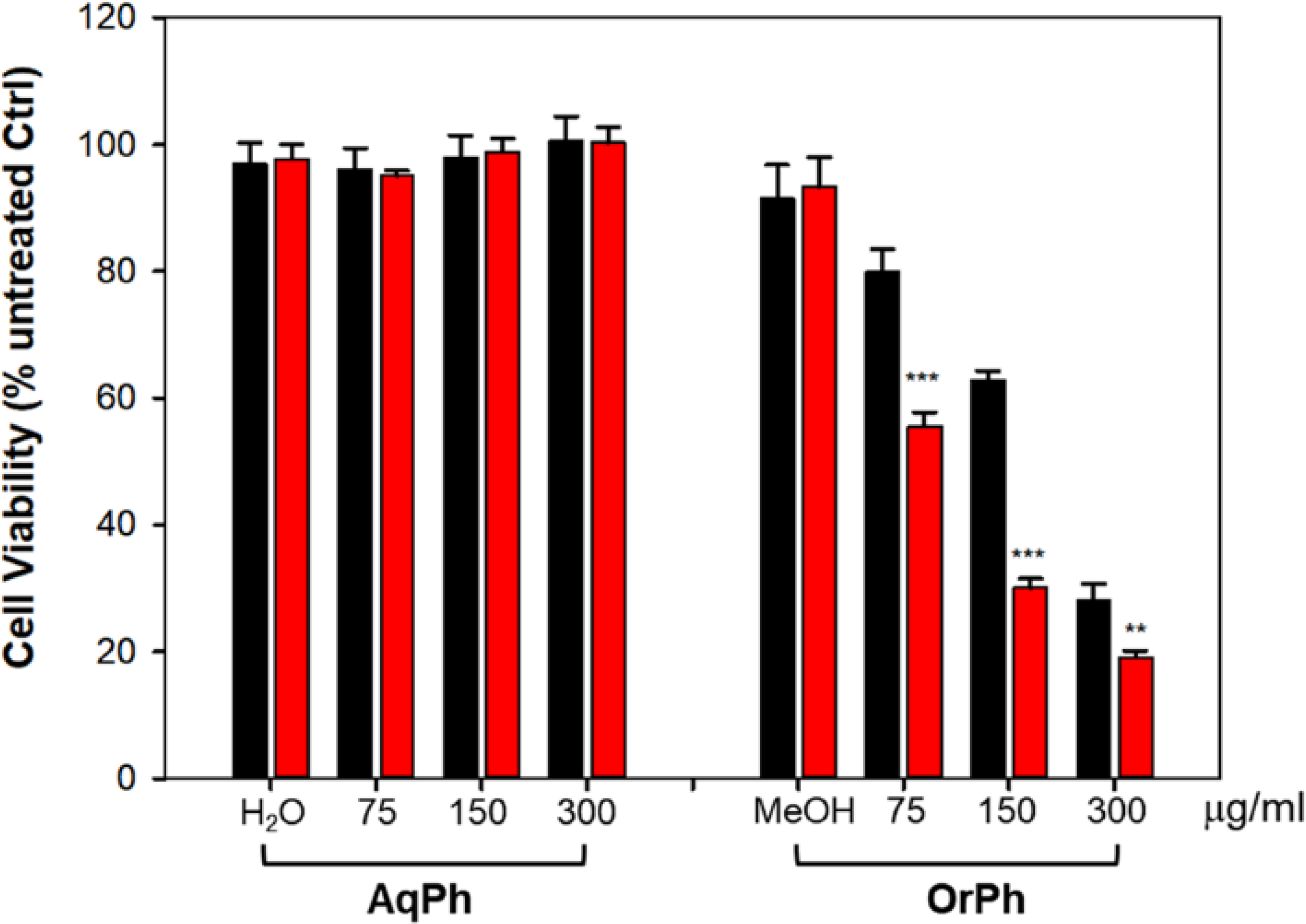
Cytotoxicity of *Ciona robusta* aqueous and organic extracts on HT-29 cell line. Cells (2×10^4^/well) were incubated with increasing concentrations (75-300 µg/ml w/v) of the aqueous (AqPh) and organic (OrPh) phases prepared as reported in the Materials and Methods section for 24 (black bars) and 48 h (red bars). Cell viability was measured using the Crystal Violet assay, as reported in the Materials and Methods section. As controls (Ctrl), water (H_2_O) and methyl alcohol (MeOH; 2% v/v, final concentration) were used in the AqPh and OrPh groups, respectively. Data are presented as means of three independent experiments ± SD. Symbols indicate significance: ***P<0.0005; **P<0.005 vs untreated Ctrl (Student’s t-test).

The capacity of the OrPh fraction to induce cell death was corroborated by microscopy analyses (Figure 4). Figure 4(a) reports the morphology of HT-29 cells photographed at 48 h after treatment with the OrPh and subsequently fixed and stained with Crystal Violet. In the right panel, it is clear that the number of stained (and viable) cells is drastically reduced compared to the control experiment (left panel). Furthermore, the cytoplasm appears strongly vacuolated (red arrows). From the microscopy analysis, other information was obtained on the capacity of the OrPh fraction to induce apoptotic cell death. In fact, the very sensitive CyQuant fluorescence staining protocol allows the dye to enter the cells and intercalate into the DNA, emitting green fluorescence at 530 nm (FITC) in leaving cells. The result of this experiment is illustrated in Figure 4(b), where the appearance of apoptotic bodies is clearly detectable (red arrows).

**Figure 4.**
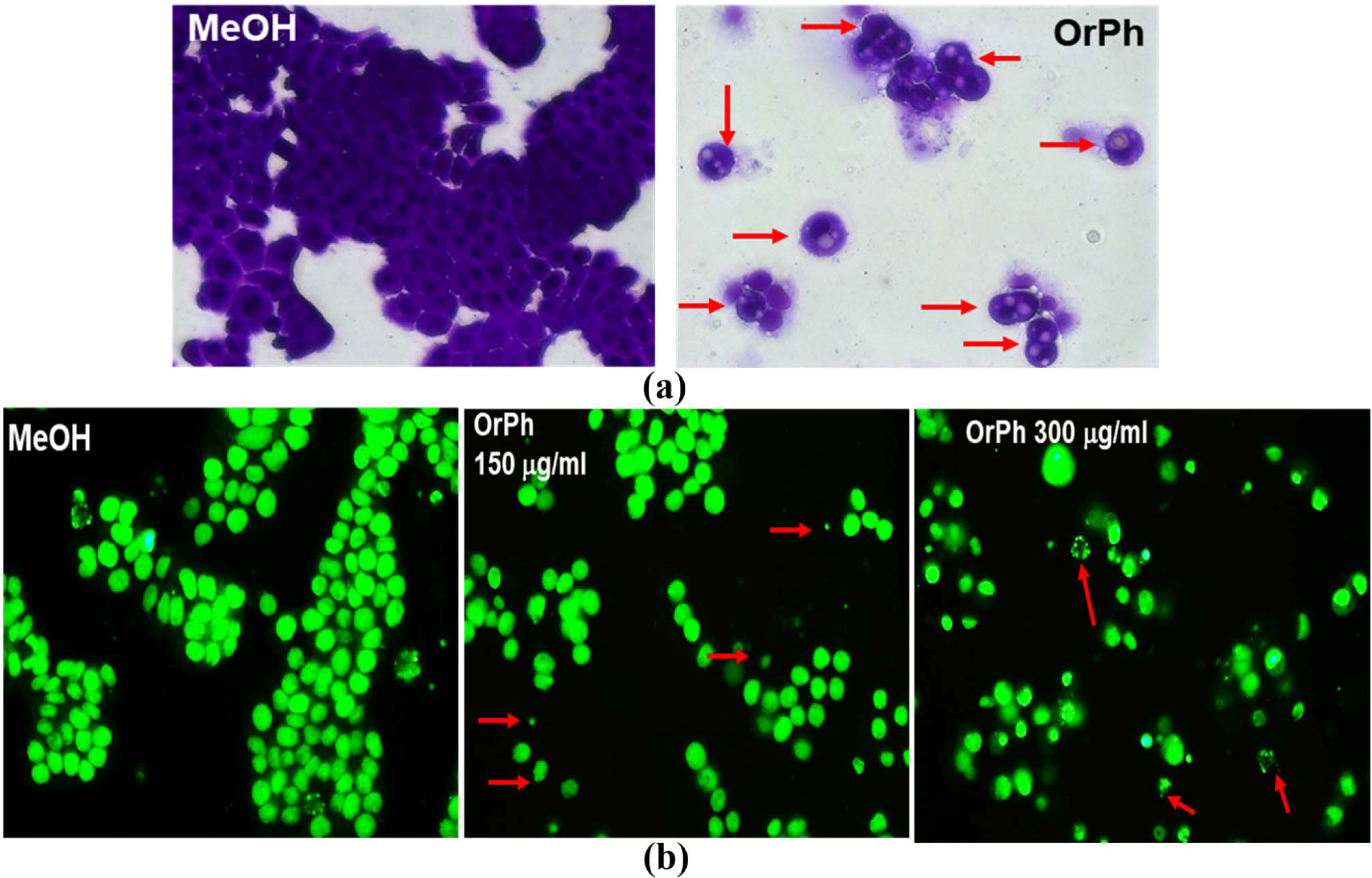
Cytotoxicity of *Ciona robusta* organic phase extract on HT-29 cell line. Panel (a): representative images (400X magnification in bright field) of HT-29 cells treated as reported in the legend of Figure 3 and stained for 48 h with Crystal Violet dye at the OrPh concentration of 300 µg/ml (w/v; right panel). The left panel indicates the treatment with the vehicle solvent (methyl alcohol; MeOH). Red arrows in the right panel indicate cells with vacuoles. Panel (b): representative images of HT-29 cells treated as reported in panel (a) at the displayed concentration of OrPh (middle and right panels) or vehicle (MeOH; left panel) and stained with CyQuant dye as reported in Materials and Methods section to evidence cell nuclei. Significant fields have been visualized with a Zeiss AXIOVERT 200 fluorescence invertoscope and photographed with a FITC filter and 400X magnification. Arrows in red indicate cells presenting apoptotic bodies.

These data confirm the original observation previously reported by our group on a similar extract prepared from ascidian *C. intestinalis* (Russo et al., 2008) where the activation of apoptosis was demonstrated by the DNA ladder profile and by the activation of caspase-3 proteolytic enzyme. However, in that dated paper, the apoptotic effects were tested on leukemia cells growing in suspension, i.e. HL-60 and HPB-ALL cell lines, which, notoriously, are more sensitive to the cytotoxic action of naturally occurring agents/extracts compared to adherent cancer cells. In fact, we also measured a strong cytotoxic activity of the OrPh fraction on HL-60 and HPB-ALL cell lines (Russo and Gallo, 2024). However, here, we demonstrated that apoptosis is also triggered by OrPh fraction in HT-29 cells, which are highly resistant to cell death induced by chemotherapy drugs, ionizing radiation (Russo et al., 2021) and apoptotic inducers, such as death receptors (Fas, TRAIL) and BH3 mimetic agents (Okumura et al., 2008; Ahmed et al., 2016; Nahacka et al., 2018).

One of the key questions associated with the antiproliferative capacity of new agents on cancer cells is the effects on normal cells to assess their specificity for the malignant phenotype and provide information on the risks vs benefits ratio. The advantage of using HT-29 cell line is represented by the capacity of these cells to differentiate when treated with differentiating agents such as sodium butyrate (NaBt) assessed by several biochemical and morphological markers. In fact, NaBt increases the activity of alkaline phosphatase enzyme (AP), stimulates the appearance of mucin-producing cells, the formation of highly differentiated goblet-like enterocytes, and induces the expression of differentiation-associated cytokeratin proteins (Tanaka et al., 1990; Russo et al., 1997). These changes, mimicking the phenotype of normal intestinal epithelial cells, support the use of NaBt-differentiated HT-29 cells as a suitable model to compare the cytotoxicity of a given agent/extract in malignant cells *vs* their differentiated and normal-like counterpart.

On these premises, HT-29 cells were induced to differentiate with 4 mM NaBt, as demonstrated by the increase of about 9-fold of the AP activity (Figure 5(a)) between 96 and 120 h from the treatment. Differentiated cells were treated with increasing concentrations of OrPh fraction in the range of 75-300 mg/ml for 24 h. As reported in Figure 5(b), the cytotoxic effect of the extract was minimal compared to undifferentiated HT-29 cells treated with identical concentration and time (Figure 3; bars on the right part of the graph).

**Figure 5.**
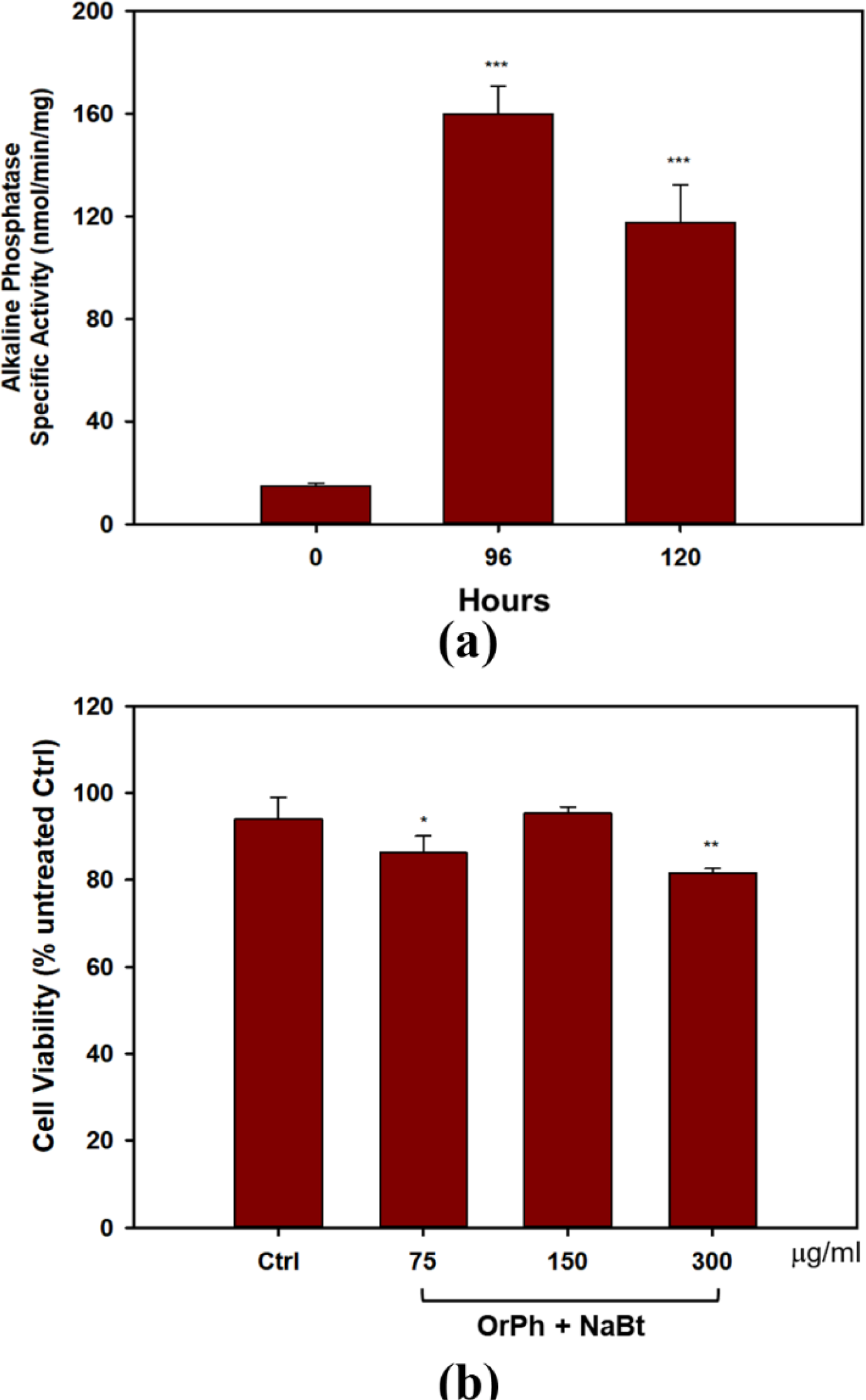
Effect of *Ciona robusta* organic phase extract on differentiated HT-29 cell line. Panel (a): Cells (2×10^4^/well) were incubated with 4 mM NaBt for the indicated times, harvested and lysed. The extracts were assayed to measure the activity of AP enzyme expressed as nmoles of pNPP (p-nitrophenyl phosphate) per minute per mg of total proteins assayed as reported in the Materials and Methods section. Data are presented as means of two independent experiments ± SD. Symbols indicate significance: ***P<0.0005 vs NaBt treated cells at time 0 (Student’s t-test). Panel (b): Cells treated with NaBt as described in panel (a) were added with the indicated concentrations of OrPh for 48 h. Cell viability was determined using the Crystal Violet assay, as reported in the Materials and Methods section. Control (Ctrl) point is represented by cells treated with MeOH (2% v/v). Data are presented as means of two independent experiments ± SD. Symbols indicate significance: **P<0.005; *P<0.05 *vs* Ctrl (Student’s t-test).

This experiment is promising in supporting the hypothesis that the toxicity of the active component(s) present in the *C. robusta* extract is specific for the malignant phenotype and does not interfere with the physiology of differentiated cells. It is important to remember that differentiated cells do not divide and are normally arrested in the G1 phase of the cell cycle. Therefore, the massive cell death observed in undifferentiated HT-29 treated with the extract suggests that the bioactive compound(s) may trigger key modulators of the pathways regulating the G1/S transition or the G2/M phase of the cell division cycle.

Of course, data reported in Figure 5, although encouraging, should be considered only preliminary in assessing the absence and/or very limited cytotoxicity of *C. robusta* extract on normal cells. Screening other cell types is currently under scrutiny (e.g., human peripheral blood mononuclear cells from healthy donors, primary cells, etc.).

When working with reactive extracts possessing biological activities, it is necessary to exclude artifactual phenomena due to the possibility that the observed cytotoxicity could be associated with the generation of hydrogen peroxide and/or other reactive oxygen species (ROS) following the interaction between the extracts and the cell culture medium components. This possibility has been well described in the literature (Halliwell and Whiteman, 2004; Halliwell, 2009). To this aim, we first measured the formation of H_2_O_2_ by FOX assay after incubation of OrPh and AqPh extracts at the concentrations of 500 µg/ml (w/v) with cell culture medium in the absence of cells. The FOX method is widely used to measure levels of hydrogen peroxide and more generally hydroperoxides in biological systems. As an example, the method is used for the detection of levels of lipid hydroperoxides in plant, fruit and vegetable tissues (DeLong et al., 2002). The results presented in Figure 6(a) indicated that while the OrPh fraction generated a significant amount of H_2_O_2_ (about 85 µM), a minimal production was observed for the AqPh fraction (about 10 µM). However, when we exposed HT-29 cells to increasing concentrations of H_2_O_2_ (up to 1000 µM), no cytotoxicity was observed (Figure 6(b); left side of the graph), compared to about 65% of cell death caused by the OrPh fraction alone (Figure 6(b); right side of the graph).

**Figure 6.**
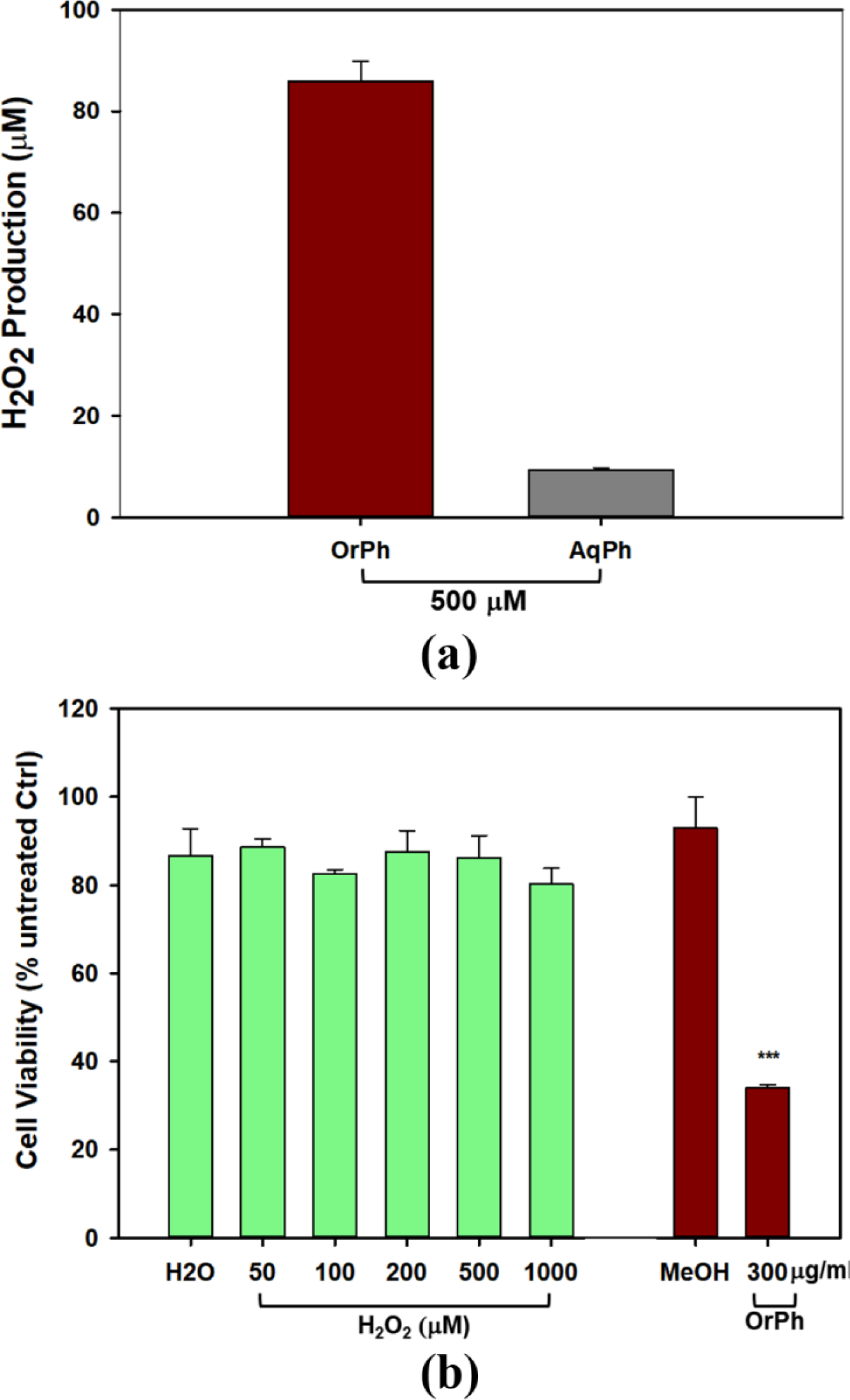
Effect of hydrogen peroxide production by *Ciona robusta* aqueous and organic extracts on HT-29 cell line. Panel (a): the FOX assay was performed starting from the indicated concentration (500 µg/ml w/v) of both the aqueous (AqPh) and organic (OrPh) phase extracts in the absence of cells as reported in Materials and Methods section. The production of H_2_O_2_ was measured after 24 h of incubation of the cell culture medium in the presence of the extracts. Panel (b): Cells (2×10^4^/well) were incubated with increasing concentrations (50-1000 µM of H2O2 (green, left bars) or OrPh extract (300 µg/ml w/v) (dark right bars) for 24 h and cell viability was measured using the Crystal Violet assay as reported in Materials and Methods section. As controls, water (H_2_O) and MeOH (2% v/v) were used in the experiments, respectively. Data are presented as means of two independent experiments ± SD. Symbols indicate significance: ***P<0.0005 *vs* control (MeOH) (Student’s t-test).

The results presented in Figure 6 confirm the low response of HT-29 cells to oxidative insults, which justify their high resistance to radio- and chemotherapy (Russo et al., 2021; Russo et al., 2023). However, and even more importantly, these data support the evidence that *C. robusta* OrPh extract contains agent(s) able to interfere with the processes regulating cell growth and cell death in HT-29 that cannot be attributed to artefacts generated by the experimental conditions and are due to the direct interaction of such as putative extract components with the malignant cells. This simple experiment is of crucial importance considering the role of ROS and oxidative damage in pathology (Forman and Zhang, 2021) and the importance of their correct determination in the different experimental settings, both in cells and *in vivo* (Murphy et al., 2022). In addition, in recent years, qualified literature and numerous research groups in the field suggest caution when interpreting data on the biological properties of molecules/extracts deriving from naturally occurring sources since many of them may fall within the category known as PAINS (pan-assay interference compounds) that function as reactive chemicals rather than target-directed drugs (Russo, 2007; Baell and Walters, 2014). For example, PAINS are responsible for the unspecific production of hydrogen peroxide that causes interference with the redox regulatory mechanisms.

From the careful microscopy observation of data reported in Figure 4, it emerged that HT-29 cells treated with the OrPh extract at the highest concentrations (300 μg/ml w/v) showed the presence of vacuoles of different sizes in the cytoplasm or the fragmentation of the cytoplasm itself (Figure 4(a)). Nuclei stained with the CyQuant fluorescent dye appeared normal in size until 48 h, and apoptotic bodies were detectable at 72 h (Figure 4(b)). This latter observation suggests that the treatment with OrPh extract could activate the phenomenon of cytotoxic autophagy or type II cell death accompanied by the onset of apoptosis at a later time (Doherty and Baehrecke, 2018; Russo and Russo, 2018). To confirm this hypothesis with quantitative measurements, cells treated with different concentrations of *C. robusta* fractions for 48 h were stained with a selective fluorescent reagent (Cyto-ID assay) able to detect phagosomes, autophagosomes, and autolysosomes, which are formed in the different phases of autophagy. In contrast, the nuclei were stained with Hoechst fluorochrome 33342. As reported in Figure 7(a), autophagy vacuoles characterized by green staining (FITC) were evidenced in the treated cells, while the nuclei emitted fluorescence at a blue wavelength (DAPI). The quantification of the vacuoles was determined by spectrofluorometric measurement of the FITC/DAPI ratio (Figure 7(b)). A significant increase was observed in the number of autophagic vacuoles compared to the control at the highest concentration (300 μg/ml) applied of the OrPh fraction, but also at the lowest (75 μg/ml), where the level of cytotoxicity was not maximal.

**Figure 7.**
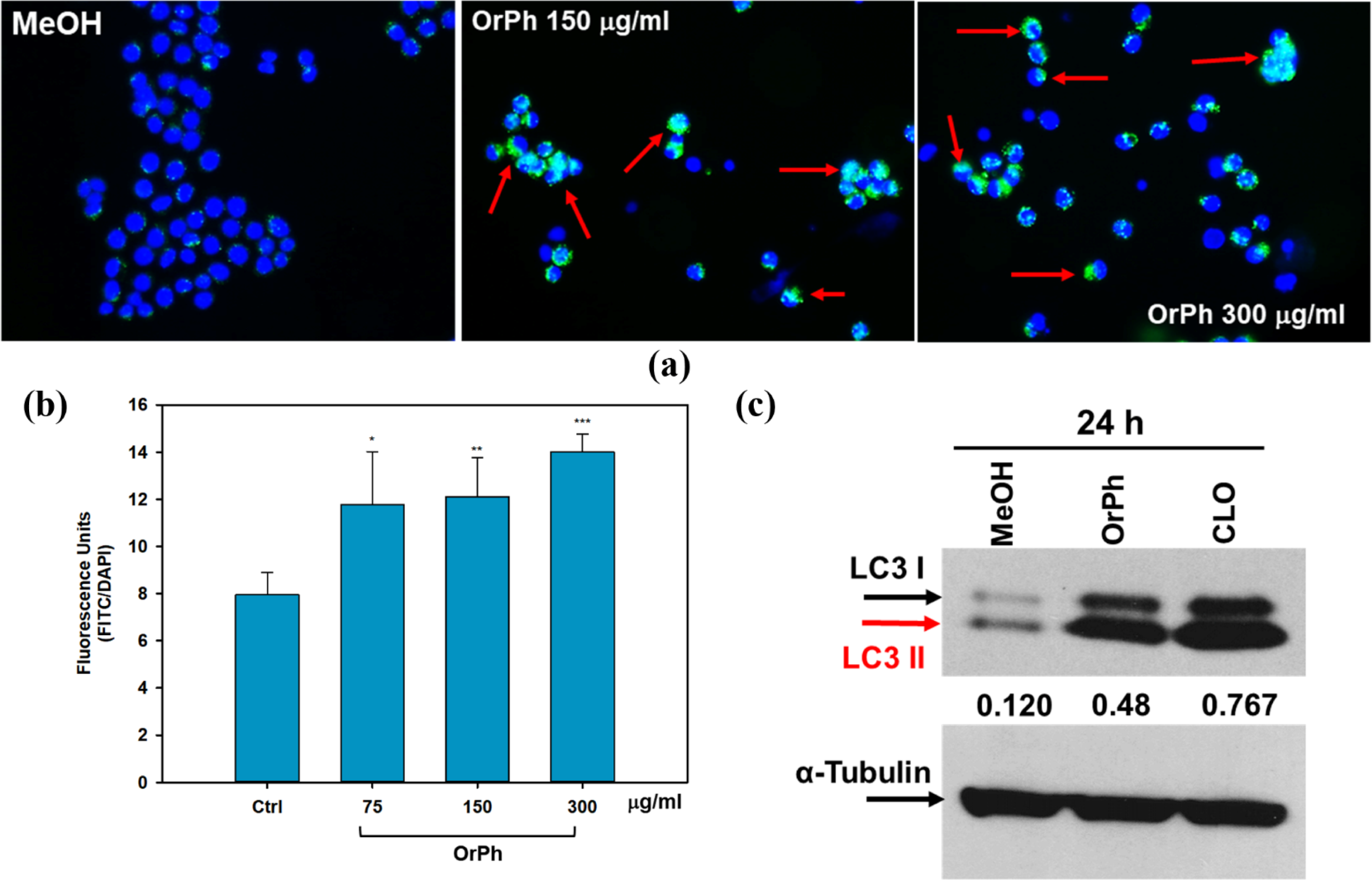
Activation of the autophagy in HT-29 cells treated with *C. robusta* organic phase extracts. Panel (a): representative images evidencing the autophagic vacuoles (in green) and the cell nuclei (in blue) of using double staining with Cyto-ID/Hoechst. HT-29 cells were treated at the indicated concentrations of OrPh fraction (middle and right panels) or vehicle (left panel) for 48 h and stained with Cyto-ID/Hoechst, as reported in the Materials and Methods section. Significant fields have been visualized with a Zeiss AXIOVERT 200 fluorescence invertoscope and photographed with a FITC filter and 400X magnification. Red arrows indicate cells with autophagosomes. Panel (b): Fluorometric quantification of autophagosomes in HT-29 cells treated as reported in panel (a) with the indicated different concentrations of OrPh fraction. The bars indicate S.D. The significance of the treatments compared to the vehicle solvent (Ctrl, methyl alcohol) was calculated with the Student’s test: *p<0.05, **p<0.005, ***p<0.0005. Panel (c): Immunoblotting to detect LC3 I-II in HT-29 cells treated with 300 mg/ml (w/v) of OrPh fraction. Chloroquine (20 µM, CLO) was used as a positive control. Densitometry values are indicated by numbers between the upper and lower panels and are calculated as the ratio between LC3-II (active isoform) and α-tubulin expressions.

To confirm the activation of the autophagic process, biochemical markers, such as the expression of LC3 isoforms, were employed. HT-29 cells were treated for 24 h with 300 µg/ml OrPh extract, and cell lysates were separated on SDS-PAGE to detect the expression of LC3-I/II by immunoblotting. The LC3 system mediates the formation of the autophagosome. It is present in two forms: the free cytosolic and inactive form (LC3-I) and the active form conjugated to phosphatidylethanolamine (LC3-II) (Klionsky et al., 2021). Figure 7(c) shows a 4-fold increase in the lipidated form of LC3 (LC3-II) after treatment with the extract compared to MeOH-treated control cells. The growth was comparable to that induced by chloroquine (CLO), used in the experiment as a positive control (Russo and Russo, 2018).

These results suggest that the OrPh extract prepared from *C. robusta* induces a change in the autophagy flux in HT-29 cells as early as 24 h after treatment. At longer times (48-72 h), a threshold is reached beyond which the autophagic process becomes lethal and the cell death process is activated (Doherty and Baehrecke, 2018). It cannot be excluded that, alongside cytotoxic autophagy, necrosis and/or apoptosis may also occur, and future data (caspase activity measurement and apoptotic body measurement, assessment of Annexin V positivity) will be able to confirm this hypothesis. In fact, our previous study (Russo et al., 2008) demonstrated that a partially purified extract from *C. intestinalis* had a cytotoxic effect on various transformed cell lines, including those of colon adenocarcinoma Caco-2, activating short-term apoptosis (caspase-3 activation at 12 h and nuclear DNA fragmentation at 24 h).

To verify whether the antiproliferative activity of the OrPh extract prepared from *C. robusta* tissues was attributable to the presence of symbiotic microorganisms, colonies of *C. robusta* consisting of several dozen specimens were collected in the Gulf of Naples, maintained in aquaculture conditions and treated with antibiotics (100 mg/L of gentamicin and kanamycin) for three weeks. Groups of control animals were kept under the same experimental conditions but in the absence of antibiotics. The OrPh extracts were prepared from antibiotics-treated and untreated samples and assayed on HT-29 cells at the above experimental conditions. The rationale of the experiment was to verify if, by removing the potentially antibiotics-sensitive microorganisms present in the adults of *C. robusta*, the cytotoxicity of the OrPh extract would be abolished. Data presented in Figure 8 clearly show that the presence of antibiotics does not modify the cytotoxic profile of the extracts, suggesting that the active compounds may not have a symbiotic origin. The reduced cell viability was even more pronounced in the antibiotics group than in the extract from control individuals. However, this difference may be due to internal variability in the preparation procedures. It is essential to underline the expected behaviour of the two OrPh extracts that were not influenced by the extensive treatment (up to three weeks) with a mixture of antibiotics. It is worthwhile to mention that maintaining individuals in microbiologically sterile conditions under the constant presence of a cocktail of antibiotics does not cause any apparent functional or physiological alteration. As an example, *in vitro* fertilization tests did not differ between the two groups concerning larval development.

**Figure 8.**
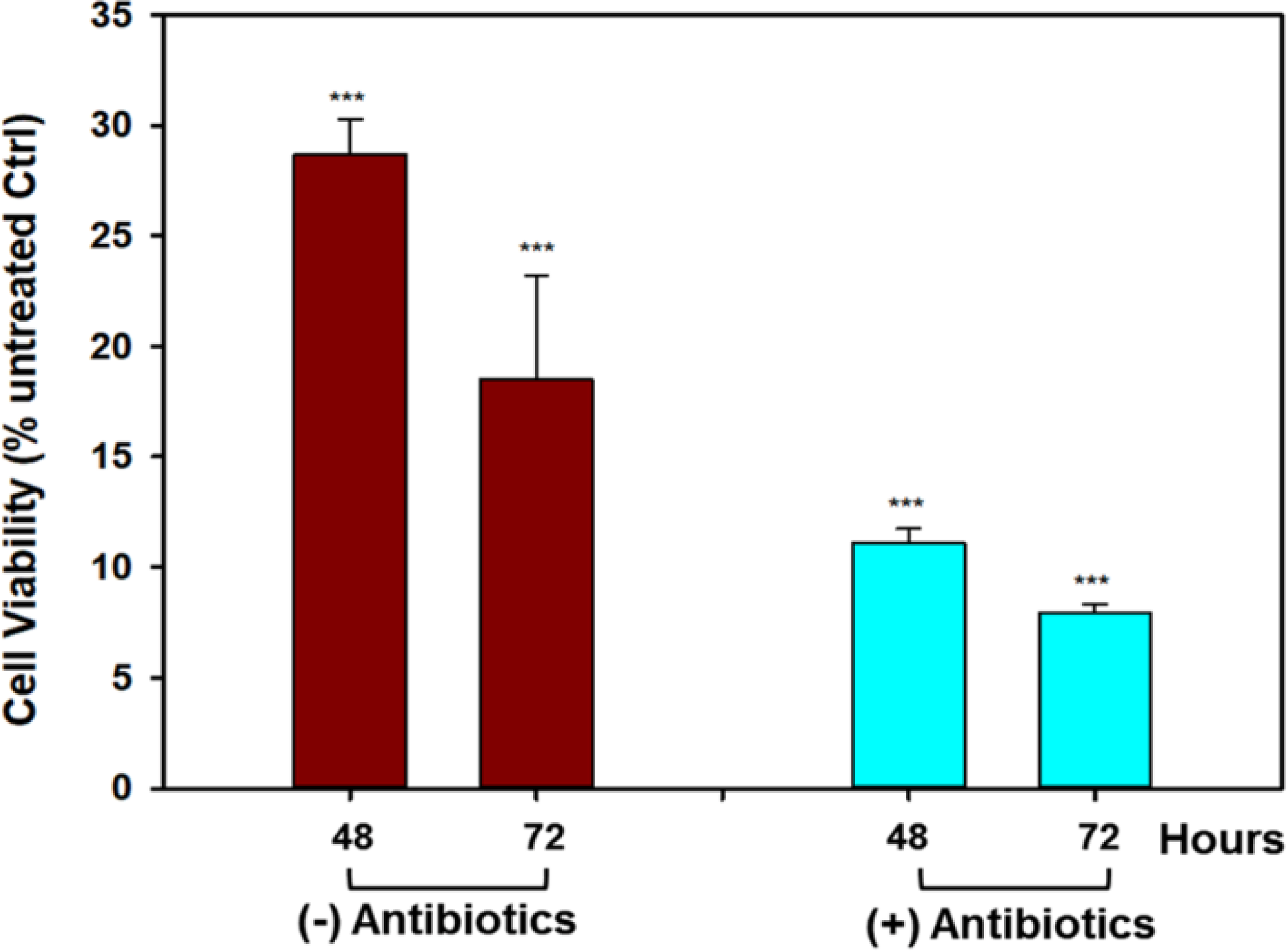
Effect of antibiotics treatment on the cytotoxicity induced by *Ciona robusta* organic extract on HT-29 cell line. Cells (2×10^4^/well) were incubated with a concentration of OrPh phase corresponding to 300 µg/ml (w/v) for the indicated times and prepared from *C. robusta* colonies treated for three weeks with a cocktail of antibiotics (100 mg/L of gentamicin and kanamycin). After antibiotics incubation, the extracts were prepared as reported in the Materials and Methods section and cell viability was measured using the Crystal Violet assay. Data are presented as means of two independent experiments ± SD. Symbols indicate significance: ***P<0.0005 *vs* cells treated with the vehicle solvent (MeOH 2% v/v; Student’s t-test).

The role of symbionts in assessing the presence of bioactive compounds with potential pharmacological properties in ascidians has been largely debated and recently extensively reviewed (Watters, 2018; Cooreman et al., 2023). It is well established the richness of the microbial diversity associated with ascidians (reviewed in (Cooreman et al., 2023)) and *C. intestinalis* represents a good example with about 33 different microbial genera, dominated by the gammaproteobacteria class, in the gut microbiota (Utermann et al., 2020). However, if from one side the structural resemblances existing between natural products isolated from marine organisms and metabolites produced by microorganisms suggest common biosynthetic origin thus supporting the symbiotic theory, it is also known that the actual synthetic origin of the majority of metabolites in ascidians and other marine organisms remains undetermined (Tianero et al., 2015).

Data presented in Figure 8 is in favour the non-symbiotic origin of the active compounds present in the OrPh extract. This evidence is supported not only by the observation that keeping the ascidians under sterile conditions (presence of antibiotics) does not eliminate the cytotoxic effects of the extract but also by other circumstantial evidence: i. OrPh extracts prepared from individuals collected from different geographic areas and latitudes, such as the Mediterranean Sea (Gulf of Naples and Gulf of Taranto; Italy) and Atlantic Ocean (Roscoff; France), where it is unlike the microbial communities are identical, show comparable cytotoxic activity (Russo and Gallo, 2024); ii. Similar antibiotic treatments have been applied to different marine invertebrates to detect the presence of symbiotic microorganisms and their contribution to the biosynthesis of biologically active secondary metabolites (Davidson et al., 2001; Dedeine et al., 2001; Hildebrand et al., 2004); iii. Examples exist in the literature of marine organisms that, after pre-treatment with antibiotics, could still biosynthesize the bioactive compounds (Hildebrand et al., 2004).

## 4 Conclusions

The first important novelty of the present work concerns in the demonstration that even in a Mediterranean specie of tunicates, i.e. *Ciona robusta*, bioactive compounds with anti-proliferative activity are present. Furthermore, since the procedure followed for the preparation of the organic extract closely resemble the one applied to the isolation of ET-743 (Ecteinascidin-743; trabectedin), one of the most clinically successful anti-tumor molecule isolated from the ascidians and approved by the FDA for clinical use under the brand name Yondelis®, we hypothesize that the active compound(s) present in the *C. robusta* organic extract may be structurally similar to ET-743. We are aware that one limitation of the present study regards the absence of information on the chemical characterization of the bioactive extract. We are intensively working on the isolation and identification of the antiproliferative component(s), a process that is time-consuming that requires large amounts of tissue/animals considering that the recovery yield of the bioactive compounds is extremely low (< 1%).

The present work also investigated the “symbiotic” origin of the bioactive compounds present in the *C. robusta* extract. It is known that marine invertebrates host in their tissues a multitude of microorganisms, including bacteria, cyanobacteria and fungi that reside in the extra- and intra-cellular space and, in some cases, these symbiotic microorganisms can represent up to 40% of the entire biomass (Proksch et al., 2002). As discussed above, genetic analysis has highlighted that many bioactive peptides are synthesized by obligate symbiotic microorganisms not from the parent organism, as for ET-743. For this reason, many authors have hypothesized that most bioactive compounds isolated from marine invertebrates derive from symbiotic bacteria/microalgae or are assimilated by the animal through the food chain (Proksch et al., 2002). However, in our case, ascidians maintained “in sterility” did not show any apparent signs of functional abnormality (e.g., *in vitro* fertilization tests indicated no differences between antibiotics-treated and -untreated groups concerning larval development). We presented data suggesting that the antibiotic treatment does not modify extract’s anti-proliferative activity.

Finally, the most important novelty of the present work has been the demonstration that, at least in HT-29 cells, the organic extract induced cytotoxic autophagy. In 2016, Ruocco et al. (Ruocco et al., 2016) discussed blueprint autophagy, so defined because it is induced and/or inhibited by natural products of the navy. However, in this article, none of the molecules reviewed were derived from tunicates. Therefore, to our knowledge, the data reported in the present work represent one of the first evidence of autophagic activity induced by compounds derived from sea squirts.

In conclusion, we demonstrated the presence of molecules not of symbiont origin in the tissues of *C. robusta* and the autophagic activity of these compounds. Therefore, *C. robusta* can be proposed as a new model for studying the mechanisms of autophagy since essential genes regulating the autophagic process have been identified in the genome of this organism (Godefroy et al., 2009). Therefore, the presence of potential compounds capable of modulating the autophagic response could not only have a possible use in tumor therapy, as for other compounds derived from marine organisms but also perform essential physiological functions for understanding the regulation of chordate development during the evolution.

## 6 Author Contributions

GLR, ET and AG: Conceptualization. MR, MR, YMP and AG: methodology, validation and investigation. GLR: writing-original draft. AG and YMP: writing-review and editing. YMP: visualization. GLR, and ET: supervision. GLR, ET and AG: funding acquisition. All authors have read and agreed to the published version of the manuscript.”

## Acknowledgements

The authors thank Mr. Alberto Macina of the Stazione Zoologica Anton Dohrn for technical support in ascidian’s maintenance.

## Footnotes

1 https://www.ncbi.nlm.nih.gov/datasets/genome/?taxon=7713

